# Early alterations of thalamo- and hippocampo-cortical functional connectivity are biomarkers of epileptogenesis after traumatic brain injury

**DOI:** 10.1101/2023.03.08.531764

**Authors:** Marina Weiler, Evan S. Lutkenhoff, Brunno M. de Campos, Raphael F. Casseb, Paul M. Vespa, Martin M. Monti, the EpiBioS4Rx Study Group

**Author notes:** Correspondence to: Martin M. Monti, Department of Psychology, University of California Los Angeles, Los Angeles, CA. 90095, USA.

## Abstract

The Epilepsy Bioinformatics Study for Antiepileptogenic Therapy (EpiBioS4Rx) study is a prospective multicenter clinical observational study to identify early biomarkers of epileptogenesis after moderate-to-severe traumatic brain injury (TBI). In this preliminary analysis of 37 patients, using a seed-based approach applied to acute (i.e., ≤ 14 days) functional magnetic resonance (MRI) imaging data, we directly test the hypothesis that the epileptogenic process following brain trauma is associated with functional changes within hippocampal and thalamo-cortical networks. Additionally, we hypothesize that the network connectivity involving thalamic and hippocampal circuits underlying early and late-onset epileptogenesis would differ. The three groups did not differ by sex distribution (*χ*^2^_(2)_ = 1.8, *p* = .407), age (*H*_(2)_ = 4.227, *p* = .121), admission Glasgow Coma Scale (*H*_(2)_ = 3.850, *p* = .146) or postinjury day of the MRI session (*H*_(2)_ = .695, *p* = .706). The primary finding is that patients with early seizures, a sign of early epileptogenesis, exhibited pattern 1, namely, an increased positive connectivity in thalamic and hippocampal networks, as compared to patients who had no epileptogenesis, or late epileptogenesis (*p* < .05, FWE-corrected at the cluster level). In contrast, this finding was absent in those patients who exhibited late seizures, with the latter group displayed pattern 2, namely, a lower positive and higher negative connectivity in the hippocampal network, as compared to patients who had no signs of epileptogenesis (*p* < .05, FWE-corrected at the cluster level). Patients with either pattern 1 or pattern 2 connectivity profiles in thalamic and hippocampal networks were significantly predictive of late (*i.e*., between 7 days and 2 years) epileptogenesis following brain trauma. A Receiver Operating Characteristic (ROC) Curve analysis model that included thalamic and hippocampal functional connectivity values presented an Area Under the Curve (AUC) 87.7, specificity 86.7, and sensitivity 84.6. Our results indicate that dysfunction in hippocampal and thalamo-cortical networks are potential biomarkers for early and late epileptogenesis following a TBI.

## INTRODUCTION

Posttraumatic epilepsy (PTE) is a common long-term health problem associated with traumatic brain injury (TBI) (Lowenstein, 2009), as well as a risk factor for long-term mortality after TBI (Uski et al., 2018). Increased incidence of seizures after TBI is related to injury severity and specific injury characteristics, such as penetrating injury, skull fracture, dural injury, hemorrhagic lesion, surgical treatment, and prolonged impaired consciousness (Asikainen et al., 1999; Frey, 2003; Temkin, 2003; Xu et al., 2017). The timing of seizures after TBI is variable and defined as early (within 7 days) and late (after 7 days) post-injury. The process of epileptogenesis occurs over the latent period and it remains unclear if there are viable biomarkers of this process (Agrawal et al., 2006; Gupta et al., 2014; Pitkänen & Immonen, 2014). However, the latency period represents an exceptional opportunity to discover novel biomarkers, including non-invasive imaging biomarkers, which capture or identify the epileptogenesis process early following TBI.

With this background in mind, the Epilepsy Bioinformatics Study for Antiepileptogenic Therapy (EpiBioS4Rx) is a prospective observational study of moderate-to-severe TBI patients to identify biomarkers that may help identify and prevent seizure occurrence and better understand the mechanisms underlying PTE (Vespa et al., 2019). In this study, patients undergo continuous electrocephalography, MRI, and other serum collection starting in the first week after TBI and are followed up to 2 years. Giving the extensive literature linking seizures with abnormalities in hippocampus and thalamus in animals models of TBI (Golub & Reddy, 2022a; Immonen et al., 2013; Pitkänen et al., 2009; Shultz et al., 2013; Vespa et al., 2010), and TBI patients (Lutkenhoff, Shrestha, et al., 2020; Vespa et al., 2010), the EpiBioS4Rx project *a priori* hypothesized that acute abnormalities within hippocampal or thalamo-cortical networks could indicate the presence of an epileptogenic process in patients after moderate-to-severe TBI (Vespa et al., 2019).

Here, using a seed-based approach in acute post-injury functional magnetic resonance imaging (fMRI) data, we directly tested the hypothesis that the early epileptogenic process following TBI is associated with functional changes within hippocampal and thalamo-cortical networks. Additionally, we hypothesized that the pathological phenotypes—as reflected in differences in thalamic and hippocampal networks connectivity, underlying the emergence of early versus late epileptogenesis after brain trauma would diverge, as previously shown to occur using anatomical data (Lutkenhoff, Shrestha, et al., 2020).

## METHODS

### Patient screening and enrollment

Patients admitted into the ICU after an acute moderate-severe TBI involving a frontal and/or temporal lobe hemorrhagic contusion were screened across 12 sites. Patients were eligible for enrollment in EpiBioS4Rx project up to 72 hours post TBI. Criteria included ages 6 to 100 and Glasgow Coma Scale (GCS, Teasdale & Jennett, 1974) 3 to 13. Patients were excluded for isolated diffuse axonal injury, isolated epidural or subdural hemorrhages, isolated anoxic brain injury, pregnancy, incarceration, and pre-existing neurodegenerative or epileptic disorders (Vespa et al., 2019). Informed consent was obtained from a surrogate family member or legally authorized representative, using IRB-approved consent methods.

Longitudinal assessment for PTE was obtained at discharge and on days 30, 90, 180 days, and 1- and 2-years post-injury with the Ottman PTE Questionnaire (Ottman et al., 2010). Patients were divided into three groups: patients who experienced no seizures (no epileptogenesis group, NE), patients who experienced at least one seizure starting during the first week post-injury (early epileptogenesis group, EE) and patients who experienced at least one seizure starting after the first week post-injury, up to two years (late epileptogenesis group, LE)(Lowenstein, 2009).

Patients received 24 h cEEG for 72 h minimum during the first 7 days after TBI. Scalp cEEG monitoring was performed at the patient bedside using a 16–21 channel bipolar and referential composite montage (implemented according to each center’s intensive care unit protocols). Mandatory parameters included low frequency filter at 0.1 Hz, high frequency filter at 50 Hz, Notch Filter, and a 200 Hz minimum sampling rate. Each site used their own standard of care electrodes, including disk or needle scalp electrodes.

### Data acquisition

High-resolution MRIs were acquired on 1.5 or 3T MR systems, including anatomical (T1-weighted) and functional (T2*-weighted echo planar images) acquisitions (See **Suppl. Tables 1** and **2** for detailed parameter listing), performed up to 14 days (+4 days) post-injury.

### Data processing

Processing of the fMRI data was performed using UF^2^C (http://www.lni.hc.unicamp.br/app/uf2c, https://www.nitrc.org/projects/uf2c) (de Campos et al., 2020), a toolbox that runs in the MATLAB platform (MATLAB 2020a, The MathWorks, Inc., Natick, Massachusetts, United States) with SPM12 (http://fil.ion.ucl.ac.uk/spm/). We performed the T1-weighted image coregistration with the fMRI mean image, tissue segmentation, and spatial normalization to MNI-152 template. We preprocessed the functional images based on volumes realignment, normalization to MNI-152 template, and smoothing at 6 × 6 × 6 mm^3^ FWHM. To remove variance attributable to head and respiration/cardiac-induced motion, we applied a pipeline previously shown to present the best performance in denoising TBI fMRI datasets (Weiler et al., 2021), which consists of regressing the six head motion parameters, the five components with greater eigenvalue for cerebrospinal fluid and white matter masks (aCompCor, Behzadi et al., 2007), and any volume containing excessive motion (*i.e*., framewise displacement, FD) greater than 0.25 mm (Satterthwaite et al., 2013). Furthermore, participants were excluded if they presented less than four minutes of preprocessed data, mean FD greater than 0.25 mm, more than 20% of the volumes with FD greater than 0.2 mm, or any volume with FD greater than 5mm (Satterthwaite et al., 2013). Additional preprocessing steps included detrending, band-pass filtering (0.008–0.1 Hz), and grey matter masking of functional images.

After preprocessing, we performed a voxel-wise seed-based functional connectivity analysis using the bilateral thalamus (903 voxels, 7224 mm^3^) and hippocampus (1175 voxels, 9400 mm^3^) as seeds (MNI masks). Positive and negative correlation maps were estimated using the Pearson’s correlation coefficient, and then transformed to Fisher’s Z estimates using Fisher’s r-to-z transformation for subsequent harmonization of the data and statistical analysis.

Previous studies have reported the need to perform data harmonization in multisite MRI investigations (Chen et al., 2014). We first performed an exploratory analysis to confirm the existence of site effects in the data. Then, we controlled for site effects using the MATLAB version of ComBat v1.0.1 (Fortin et al., 2018; Fortin et al., 2017), https://github.com/Jfortin1/ComBatHarmonization), a software that removes inter-site technical variability even in small samples while preserving inter-site biological variability using a popular batch-effect correction tool used in genomics (Johnson et al., 2006). The Fisher’s-z-transformed positive and negative functional connectivity maps were harmonized across sites, while disease group, age, injury severity (*i.e*., admission GCS), and postinjury day of the MRI session were biological covariates kept during the removal of site effects.

### Data analysis

To investigate if thalamic and hippocampal functional connectivity profiles could indicate the presence of an epileptogenic process in patients after moderate-severe TBI, we performed a voxel-wise analysis of variance with the harmonized positive and negative functional connectivity maps separately as dependent variables, and group (NE, EE, LE) as the independent variable, regressing out the effects of age, injury severity, and postinjury day of the MRI session. Additionally, if the patient received any medication that could potentially affect the BOLD signal during the MRI (*i.e*., anti-seizure, anesthetic, benzodiazepine, anti-psychotic, or opioid), a separate regressor was generated containing a value of 1 at that variable, and 0 at all others. Group comparison was voxel-wise thresholded at *p* < 0.001 (uncorrected) and cluster-wise corrected using family wise error (FWE) at *p* < 0.05 using SPM12.

In addition, to assess the relative importance of thalamic and hippocampal networks in predicting vulnerability to seizures, we combined demographic, clinical, and MRI data in logistic regression models. Demographic information (age, sex, postinjury day of the MRI session), clinical data (admission GCS total), and the mean functional connectivity maps of thalamic and hippocampal networks were entered in four binomial logistic regression models to distinguish patients who did not have epileptogenesis from patients that did (collapsing the EE and LE groups), as follows: Model 1 - demographic data (age, sex, and postinjury day of the MRI session); Model 2 - demographic data and clinical data; Model 3a - demographic data, clinical data, average thalamic positive and negative functional connectivity; Model 3b - demographic data, clinical data, average hippocampal positive and negative functional connectivity; Model 4 - demographic data, clinical data, average thalamic and hippocampal positive and negative functional connectivity.

Then, to compare the relative importance of each set of variables in predicting the occurrence of late seizures, we performed a multivariate receiver-operating characteristic curve (ROC) analysis using the same four models but considering only the LE group. Given that the differences across groups mostly involved positive correlations (see **Fig 2** and **3**), we decided not to include a model containing only negative maps. All non-imaging statistical analysis was carried out using the SPSS v28 package (IBM Corp. Released 2022. IBM SPSS Statistics for Windows, Version 28.0. Armonk, NY: IBM Corp), and JASP software (Version 0.16.3, (*JASP Team*, 2022).

## RESULTS

### Demographic and clinical data

At the time of this manuscript preparation, 239 patients had been assessed for eligibility. A total of 118 patients were excluded, and 57 patients were lost to follow-up (details in **Fig 1**). Twentyseven patients were excluded after fMRI preprocessing due to preprocessing error or excessive in-scanner motion (cf., Weiler et al., 2022) rendering the data not suitable for seed-based analysis (Satterthwaite et al., 2012) (**Fig 1**). **Table 1** shows the demographic and clinical data of the 37 patients included in the final sample. The three groups did not differ by sex distribution (*χ*^2^_(2)_ = 1.8, *p* = .407), age (*H*_(2)_ = 4.227, *p* = .121), admission GCS (*H*_(2)_ = 3.850, *p* = .146) or postinjury day of the MRI session (*H*_(2)_ = .695, *p* = .706). All the patients who had epileptogenesis during the first week after TBI also presented epileptogenesis in the following 2 years.

**Fig 1:**
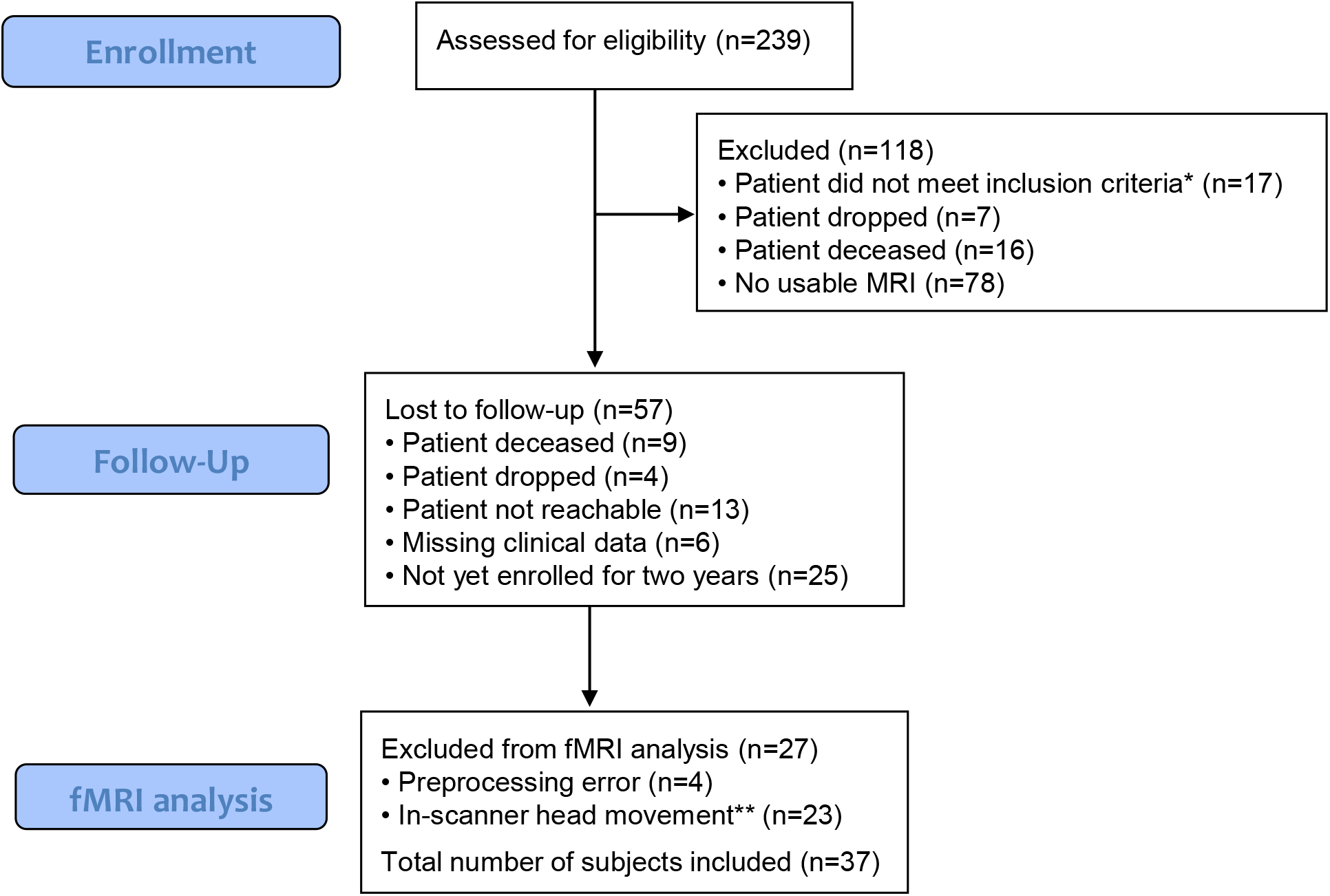
Flowchart of patient enrollment and inclusion. *Inclusion criteria according to Vespa et al., 2019. **Head movement parameters according to Weiler et al., 2022.

**Table 1:** Clinical and demographic information of patients included in this analysis.

|  | No<br>epileptogenesis | Early<br>epileptogenesis | Late<br>epileptogenesis |
| --- | --- | --- | --- |
| N | 16 | 8 | 13 |
| Age (years) | 45 ± 23 (7-84) | 47 ± 21 (19-72) | 32 ± 17 (15-65) |
| Sex (female) | 6 (37.5%) | 2 (25%) | 2 (15.3%) |
| MRI post-injury day | 9.81 ± 5.63 (0-18) | 8.38 ± 7.19 (1-17) | 10.38 ± 5.85 (1-18) |
| Admission GCS | 9.41 ± 4.25 (3-15) | 8.13 ± 4.05 (3-14) | 6.31 ± 3.37 (3-14) |
| Contusion (%) | 75 | 50 | 100 |
| Brain edema (%) | 50 | 50 | 76.92 |
| Skull fracture (%) | 93.75 | 75 | 100 |

Mean ± stdev (range). GCS = Glasgow Coma Scale.

### Harmonization of the data

To investigate and correct site effects, we assessed the z-transformed functional connectivity maps pre- and post-harmonization. We first performed an exploratory analysis to investigate any correlations between site and thalamic and hippocampal functional connectivity maps. As shown in **Suppl Fig 1**, multiple brain regions exhibited unwanted significant correlation with site (*p* < .05, FWE-corrected at the cluster-level) in the pre-harmonized images, confirming the need to perform cross-site data harmonization. These significant correlations were no longer observed after harmonization. **Suppl. Figs 2**-**7** depict the pre- and post-harmonization distributions of functional connectivity values for thalamic and hippocampal networks.

### Pattern 1: increased positive connectivity in thalamic and hippocampal networks as biomarkers for early epileptogenesis following TBI

Voxel-wise analysis revealed that the EE group presented higher positive functional connectivity in thalamic and hippocampal networks as compared to the NE and LE groups, namely pattern 1.

In EE patients compared to the NE group, the thalamus was more strongly connected to the right cuneus, and bilateral medial superior frontal regions (**Fig 2a**, **Table 2**, *p* < .05, FWE-corrected at the cluster level), whereas the hippocampus was more strongly connected to the right supplementary motor area and right cuneus (**Fig 2b**, **Table 2**, *p* < .05, FWE-corrected at the cluster level). After finding significant differences between the groups, we then explored if such difference was mainly due to a new pattern of functional connectivity in the EE group (*i.e*., an inexistent functional connectivity in the NE group between those regions), or if the functional connectivity between the regions was just stronger in the EE group. We then masked the regions that were significantly different between the groups and extracted the mean value for each patient. A one-sample t-test against zero (*i.e*., testing if that specific pattern of connectivity significantly differed from zero, meaning that the specific spatial connectivity existed within the group), was statistically significant for both groups (Thalamic network, NE group, *t_(15)_* = 9.354, *p* = < .001, 95% CI [.10, .16], Hedges’ correction = 2.21. EE group, *t_(7)_* = 11.252, *p* = < .001, 95% CI [.19, .29], Hedges’ correction = 3.53. Hippocampal network, NE group, *t_(15)_* = 10.023, *p* = < .001, 95% CI [.10, .16], Hedges’ correction = 2.38. EE group, *t_(7)_* = 17.478, *p* = < .001, 95% CI [.26, .34], Hedges’ correction = 5.49), suggesting that these patterns of connectivity are present in both NE and EE groups, but the EE group presents stronger connectivity (scatter plots in **Fig 2a** and **b**).

**Fig 2:**
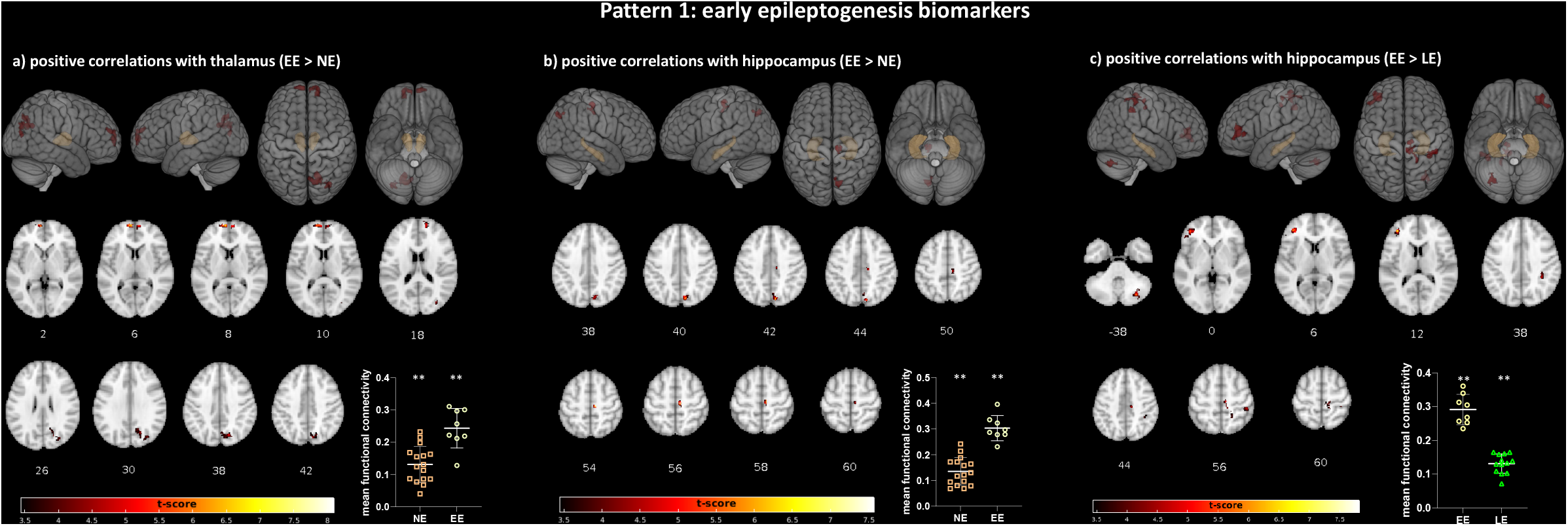
Pattern 1: early epileptogenesis biomarkers. (a) Regions presenting higher positive connection with the thalamus in the TBI early epileptogenesis group compared to the TBI no epileptogenesis group; (b) Regions presenting higher positive connection with hippocampus in TBI early epileptogenesis group compared to the TBI no epileptogenesis group; (c) Regions presenting higher positive connection with the hippocampus in the TBI early epileptogenesis group compared the TBI late epileptogenesis group (*p* < .05, FWE-corrected at the cluster level). Scatter plots showing mean functional connectivity values for the regions showing significant differences between the groups. \*\**p* < .001, one sample t-test against zero.

**Table 2:** Brain regions with higher functional connectivity in the early epileptogenesis group, compared to the no epileptogenesis group.

| Seed | Anatomical location | *p* (FWE-corr) | cluster size | T-value | Z-value | x (MNI) | y (MNI) | z (MNI) |
| --- | --- | --- | --- | --- | --- | --- | --- | --- |
| Thalamus – positive correlations | right cuneus | 0.034 | 83 | 4.7 | 3.96 | 20 | -68 | 28 |
|  | right middle occipital | 0.031 | 85 | 4.62 | 3.91 | 34 | -76 | 12 |
|  | left medial superior frontal | 0.024 | 90 | 8.1 | 5.67 | -10 | 64 | 8 |
|  | right medial superior frontal | 0.012 | 104 | 5.72 | 4.56 | 14 | 62 | 16 |
|  | right cuneus | 0.006 | 117 | 4.96 | 4.13 | 16 | -70 | 40 |
| Hippocampus – positive correlations | right supp motor area | 0.050 | 67 | 7.57 | 5.45 | 8 | -22 | 56 |
|  | right cuneus | 0.035 | 75 | 6.46 | 4.94 | 10 | -76 | 42 |

The EE group also presented higher positive functional connectivity in the hippocampal network when compared to the LE group. Regions showing stronger positive connectivity with the hippocampus included the right cerebellum, right inferior parietal, right supplementary motor area, and left middle frontal (**Fig 2c**, **Table 3**, *p* < .05, FWE-corrected at the cluster level). A one-sample t-test against zero using the connectivity values within the masked results showed that this pattern of connectivity was statistically significant for both groups (EE group, *t_(7)_* = 18.242, *p* = < .001, 95% CI [.25, .33], Hedges’ correction = 5.72; LE group, *t _(12)_* = 16.514, *p* = < .001, 95% CI [.11, .15], Hedges’ correction = 4.28), suggesting that this pattern of connectivity is present in both the EE and LE groups, but the EE group presents higher connectivity values (scatter plots in **Fig 2c**).

**Table 3:**
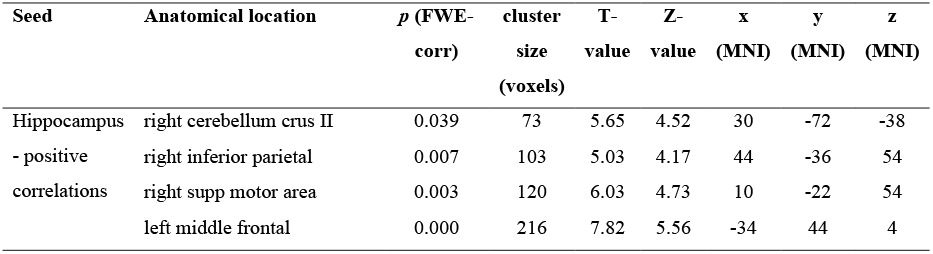
Brain regions with higher functional connectivity in the early epileptogenesis group, compared to the late epileptogenesis group.

### Pattern 2: decreased positive and increased negative connectivity in hippocampal network as biomarkers for late epileptogenesis following TBI

In addition to the lower connectivity observed in the hippocampal network in the LE group compared to the EE group described above (**Fig 2c**), a hypoconnectivity was also observed when the LE group was compared to the NE group, namely pattern 2. More specifically, voxel-wise analysis revealed that the LE group presented lower positive functional connectivity between the hippocampus and bilateral middle frontal and left cerebellum than the NE group (**Fig 3a**, **Table 4**, *p* < .05, FWE-corrected at the cluster level). A one-sample t-test against zero using the connectivity values within the masked results showed that this pattern of connectivity was statistically significant for both groups (NE group, *t_(15)_* = 10.466, *p* = < .001, 95% CI [.19, .29], Hedges’ correction = 2.48; LE group, *t _(12)_* = 13.130, *p* = < .001, 95% CI [.11, .15], Hedges’ correction = 3.41), but the LE group presented lower connectivity values (scatter plots in **Fig 3a**).

**Fig 3:**
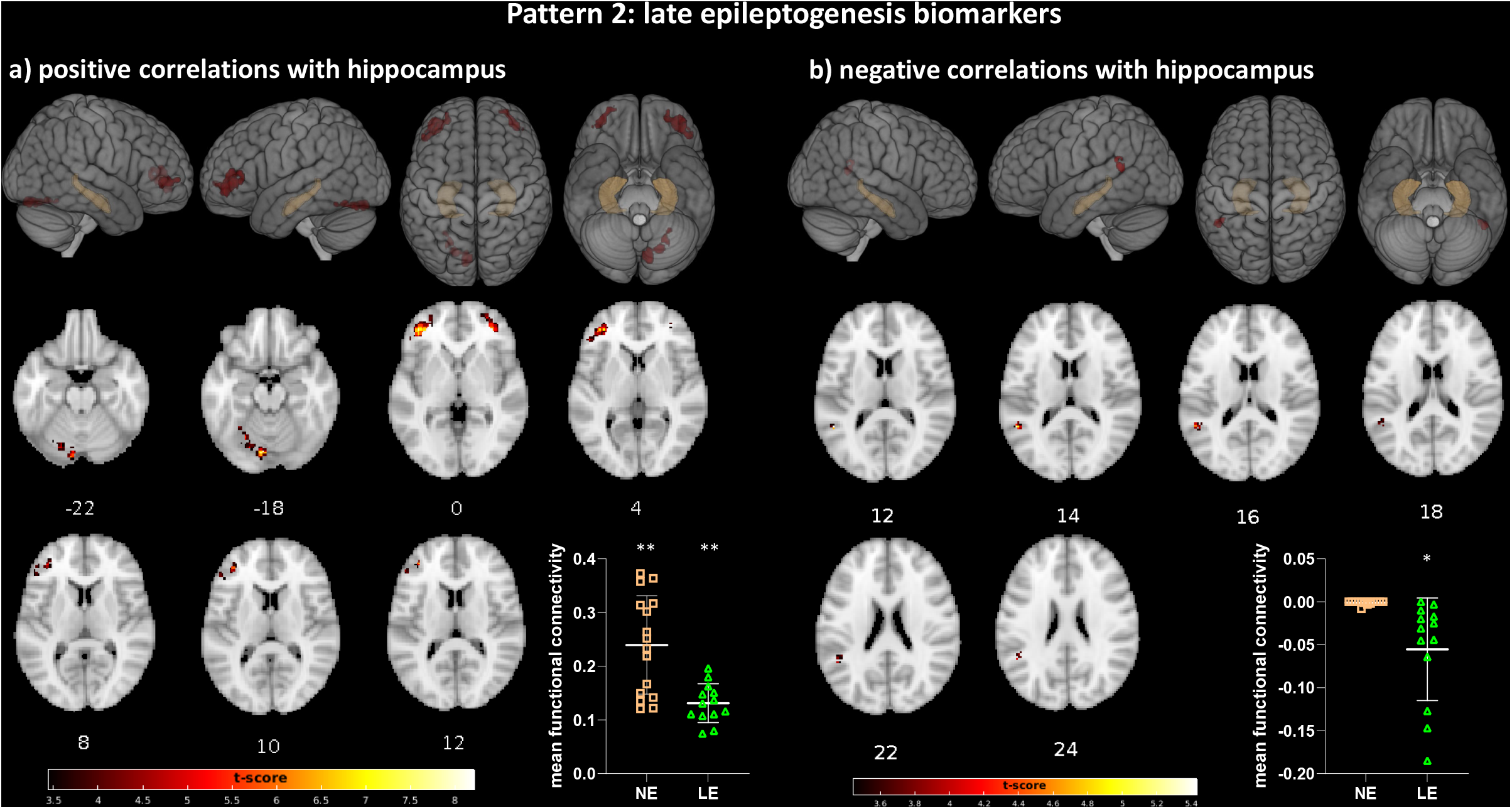
Pattern 2: late epileptogenesis biomarkers. (a) Regions presenting lower positive functional connectivity with the hippocampus in the TBI late epileptogenesis group compared to the TBI no epileptogenesis group; (b) Regions presenting increased negative functional connectivity with the hippocampus in the TBI late epileptogenesis group compared to the TBI no epileptogenesis group (*p* < .05, FWE-corrected at the cluster level). Scatter plots showing mean functional connectivity values for the regions showing significant differences between the groups. \*\**p* < .001, \**p* < .05, one sample t-test against zero.

**Table 4:**
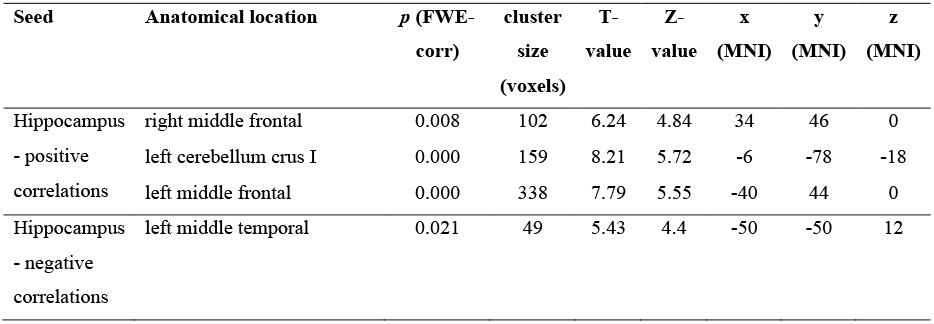
Brain regions with different functional connectivity in the late epileptogenesis group, compared to the no epileptogenesis group.

Regions presenting increased negative functional connectivity (anti-correlations) with the hippocampus in the LE group compared to the NE group included the left middle temporal region (**Fig 3b**, **Table 4**, *p* < .05, FWE-corrected at the cluster level). A one-sample t-test against zero using the connectivity values within the masked results showed that this pattern of connectivity was statistically significant for only the LE groups (NE group, *t_(15)_* = −1.668, *p* = .116, 95% CI [−.002, .0002], Hedges’ correction = −.396; LE group, *t _(12)_* = −3.335, *p* = .006, 95% CI [−.091, −.019], Hedges’ correction = −.866), suggesting that this pattern of anti-correlations is not present in the NE group, but only in the LE group (scatter plots in **Fig 3b**).

### Early dysfunction in thalamic and hippocampal networks predict the occurrence of late epileptogenesis

When we compared the NE group vs. epileptogenesis group, this latter combining EE and LE patients, all binomial logistic regression models that included demographic information, clinical data, and the mean functional connectivity maps of thalamic and hippocampal networks as predictors, failed to show statistically significant results (Model 1, *χ*^2^_(3)_ = 3.415, *p* = .332; Model 2, *χ*^2^_(3)_ = 6.502, *p* = .165; Model 3a, *χ*^2^_(3)_ = 6.718, *p* = .348; Model 3b, *χ*^2^_(3)_ = 7.432, *p* = .283; Model 4, *χ*^2^_(3)_ = 6.538, *p* = .366; and Model 5, *χ*^2^_(3)_ = 7.601, *p* = .473).

However, when we aimed to compare the relative importance of each set of variables in predicting the occurrence of late epileptogenesis (LE group vs. NE group) into the multivariate ROC analysis, models were statistically significant. Amongst the measures calculated to evaluate the models, we describe below the Nagelkerke Pseudo-*R*^2^ (a logarithmic scoring rule; the higher the score the higher the fitness of the predictive model); specificity (the percentage of TBI patients that did not have signs of epileptogenesis and were correctly predicted by the model— *i.e*., true negatives); sensitivity (the percentage of TBI patients that had signs of late epileptogenesis and were correctly predicted by the model—*i.e*., true positives); area under the curve (AUC, evaluates the model predictions using a probabilistic framework, showing the relationship between false positive and true positive rates for different probability thresholds of model predictions). As shown in **Table 5** and **Fig 4**, the model that included thalamic and hippocampal functional connectivity values presented substantially higher metrics than the models that included demographic and clinical data only. Specifically, the AUC for model 4 was 87.7, which indicates an approximate 88% chance that the neurologist will correctly distinguish a TBI patient that will have late epileptogenesis (compared to around 74% chance if only using demographic data). The specificity for model 4 was 86.7, which indicates approximately 87% chance of correctly predicting an absence of epileptogenesis following a TBI, and sensitivity was 84.6, indicating an approximate 85% chance of correctly predicting the occurrence of epileptogenesis in a 2-year period following a TBI.

**Fig 4:**
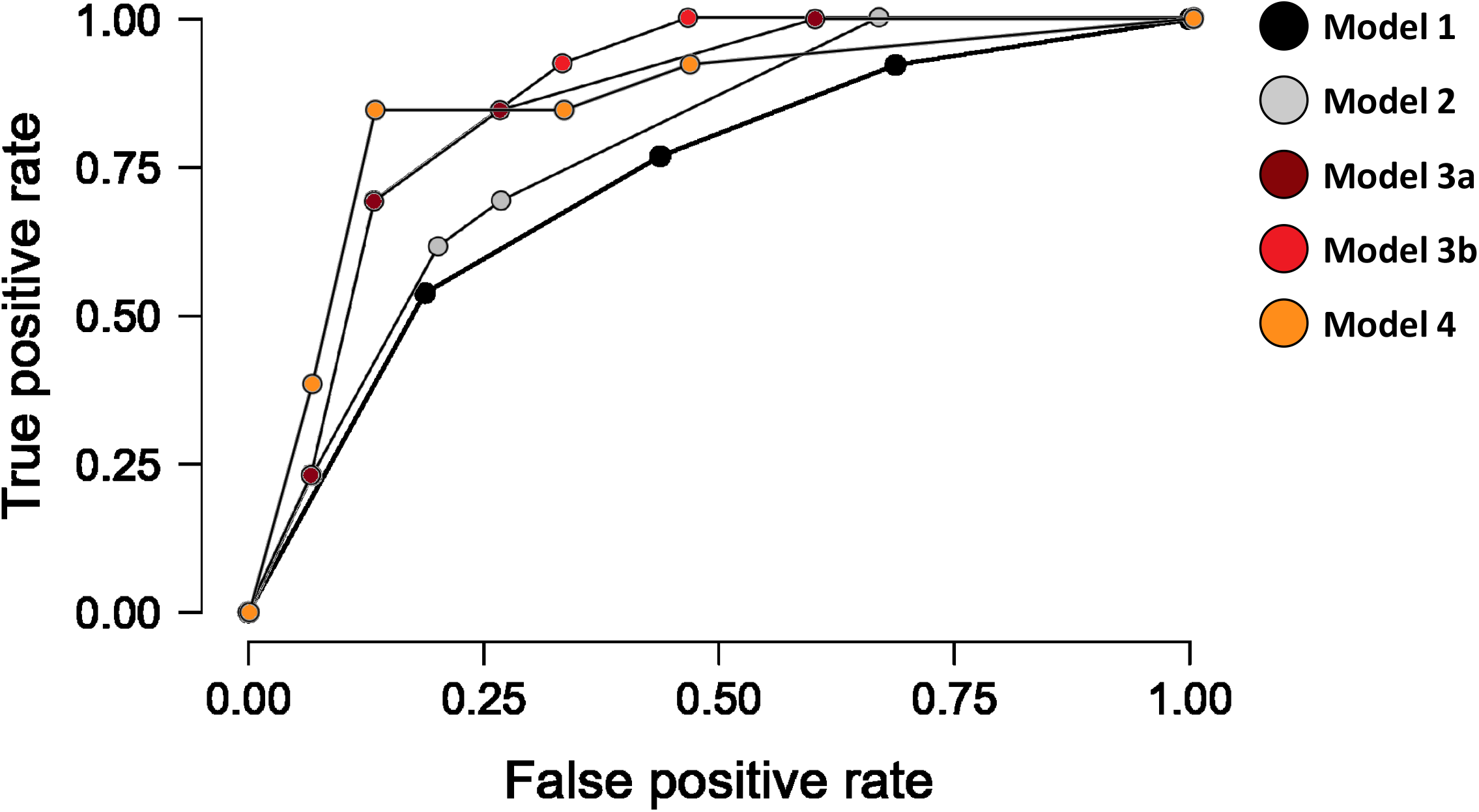
ROC plots for the four models tested to predict the occurrence of late epileptogenesis in TBI patients. Model 1: age + sex + postinjury day of the MRI session; Model 2: age + sex + postinjury day of the MRI session + injury severity (admission GCS); Model 3a: age + sex + postinjury day of the MRI session + injury severity + positive and negative thalamic functional connectivity; Model 3b: age + sex + postinjury day of the MRI session + injury severity + positive and negative hippocampal functional connectivity; Model 4: age + sex + postinjury day of the MRI session + injury severity + positive and negative thalamic and hippocampal functional connectivity.

**Table 5:** Metrics evaluating the four models used to predict the occurrence of late epileptogenesis in TBI patients.

| Model | df | $\chi^2$ | $p^*$ | Nagelkerke<br>$R^2$ | Specificity | Sensitivity | AUC |
| --- | --- | --- | --- | --- | --- | --- | --- |
| Model 1 | 25 | 4.96 | 0.17 | 0.21 | 68.8 | 61.5 | 73.6 |
| Model 2 | 23 | 8.49 | 0.07 | 0.35 | 80 | 69.2 | 79 |
| Model 3a | 21 | 11.05 | 0.08 | 0.43 | 73.3 | 76.9 | 82.6 |
| Model 3b | 21 | 12.67 | 0.04 | 0.48 | 66.7 | 76.9 | 83.1 |
| Model 4 | 19 | 15.07 | 0.05 | 0.55 | 86.7 | 84.6 | 87.7 |

\*Omnibus Tests of Model Coefficients were used to test the model fit. If the Model is significant, this shows that there is a significant improvement in fit as compared to the null model. Model 1: demographic data (age, sex, and postinjury day of the MRI session); Model 2: demographic data and clinical data; Model 3a: demographic data, clinical data, average thalamic positive and negative functional connectivity; Model 3b: demographic data, clinical data, average hippocampal positive and negative functional connectivity; Model 4: demographic data, clinical data, average thalamic and hippocampal positive and negative functional connectivity.

## DISCUSSION

Using a seed-based approach in fMRI data, we tested the hypothesis that (i) the emergence of epileptogenesis following TBI is associated with changes in brain-wide hippocampal and thalamic functional connectivity, and that (ii) connectivity changes are different for patients developing epileptogenesis shortly after injury (*i.e*., < 7 days post-injury) as compared to patients developing epileptogenesis at a later time. As hypothesized, significant differences in thalamic and hippocampal functional connectivity were observed across groups – with early and late epileptogenesis groups showing very different connectivity patterns compared with the NE group as well as each other. Specifically, we found that TBI patients presenting early epileptogenesis exhibited a hyper-positive connectivity in thalamic and hippocampal networks compared with TBI patients who presented no epileptogenesis and with TBI patients who developed late epileptogenesis (pattern 1). On the contrary, TBI patients who developed late epileptogenesis presented lower positive but higher negative connectivity in the hippocampal network when compared with the TBI group that had no epileptogenesis (pattern 2). In addition, models that included dysfunctions in thalamic and hippocampal networks were significantly able to predict late epileptogenesis following a TBI, whereas models that used only demographic and clinical data could not. Dysfunctions in thalamic and hippocampal networks were able to predict with approximately 88% chance of the occurrence of late epileptogenesis following a TBI.

So, how does our fMRI findings as discussed here link to the molecular mechanisms of epileptogenesis in unclear and yet to be fully understood. However, the pattern 1 dysfunction of the EE group could be speculated as a result of the excitotoxic environment and inflammation known to occur acutely following TBI (Lucke-Wold et al., 2015). Indeed, the imbalance between excitation and inhibition — which relates to increased extracellular glutamate in the brain and/or reduction in GABA concentrations — has been a longstanding proposed mechanism regarding ictogenesis and epileptogenesis (Huusko et al., 2015). The contribution of glutamate excitotoxicity to early onset seizure has been demonstrated in many rat models of TBI, especially in hippocampal (Drexel et al., 2015; Lowenstein et al., 1992; Zanier et al., 2003) and thalamic regions (Immonen et al., 2019; Sowers et al., 2021). In this sense, immediate and early epileptogenesis are not considered to be “epileptic” and are thought to be a direct product of the injury itself (Golub & Reddy, 2022b). This excitotoxic environment could lead to the structural alterations often observed as atrophy in TBI (Lutkenhoff, Shrestha, et al., 2020; Shultz et al., 2013; Vespa et al., 2010) which, in turn, correlate with the functional outcome of patients (Lutkenhoff, Wright, et al., 2020) and represent a risk factor for the development of PTE (Tubi et al., 2019).

Early epileptogenesis are believed to have a different underlying pathogenesis than late epileptogenesis following a TBI (Agrawal et al., 2006). Consistent with this view, we found the two groups to have different patterns of brain-wide thalamic and hippocampal connectivity, as compared with each other and with the no epileptogenesis group. The divergent profiles of pattern 1 and 2 also suggests that the neurobiological underpinnings implicated in the development of late epileptogenesis are already present acutely, well in advance of any observable behavioral or manifestation. In addition, logistic regression models aiming to predict the occurrence of any epileptogenesis following a TBI (*i.e*., combining the early and late epileptogenesis groups) failed to find statistically significant biomarkers, further corroborating the hypothesis that early and late epileptogenesis present different underlying pathogenesis. Conversely, when we differentiated the two groups, the models could significantly predict which patient experienced, over the following 24 months, late epileptogenesis. Importantly here, only the models that included hippocampal and thalamic functional connectivity data could correctly classify patients into no epileptogenesis or late epileptogenesis, whereas the models that included only demographic and clinical data could not. It should be noted that all the patients with an early epileptogenesis also presented late epileptogenesis, indicating that the occurrence of early epileptogenesis is also a strong risk factor for the development of late ones, as previously reported (Frey, 2003; Temkin, 2003). How exactly the physiopathology giving rise to early epileptogenesis plays into developing epileptogenesis in the long term, and the degree to which this group is separable from patients developing late epileptogenesis in the absence of early ictal episodes, remains to be understood.

Interestingly, in previous work conducted in our laboratory that used similar models, the authors found that injury severity, the left temporal pole, and left frontal pole significantly associated with the probability of epileptogenesis after TBI (Lutkenhoff, Shrestha, et al., 2020; Tubi et al., 2019). In their models, early and late epileptogenesis patients were combined into one group, suggesting that structural abnormalities in such regions could play a role in predicting the occurrence of epileptogenesis, either early or late. In another study, the authors used an innovative multiplex network approach to find informative complex network features to distinguish epileptogenesis-free subjects and epileptogenesis-affected subjects with an accuracy of 70% and an AUC of 76% by using T1-weighted MRI data (La Rocca et al., 2020). In the present work, functional abnormalities in hippocampal and thalamic networks could not significantly predict the occurrence of epileptogenesis or not, but they could correctly distinguish a TBI patient that will have late epileptogenesis with an approximate 88% chance, compared to around 74% chance if only demographic data was used. In addition, this model presented approximately 87% chance of correctly predicting the absence of epileptogenesis, and approximately 85% chance of correctly predicting the occurrence of epileptogenesis in a 2-year period following a TBI. In essence, our findings, combined with those previous reports, suggest that whereas both early and late epileptogenesis patients present similar anatomical abnormalities early on, they present functional alterations with different etiologies and neurobiological underpinnings that can already be detected at a short period following the injury. It is therefore a speculative yet not improbable conjecture to propose that the pattern of thalamic and hippocampal connectivities during the initial period following the TBI in the LE patients, become more similar to the pattern of EE patients, later when they develop epileptogenesis. However, longitudinal MRI acquisitions would be necessary to test this hypothesis.

In interpreting these findings, it is important to be mindful of some limitations. Above all, the small sample size could be considered a significant weakness of this study. Many patients were dropped from the study because of the high-quality control standard adopted here (such as the low threshold for in-scanner head movement), and an important development in this area for future studies is to adopt techniques that diminish as much as possible in-scanner head movement. Second, although we tried to statistically control for the effects of any medication that could potentially affect the BOLD signal during the MRI, their neural effects are yet to be unveiled.

To sum up, the delay in the emergence of chronic epileptogenesis after the initial injury represents an exceptional opportunity for intervention with antiepileptogenesis therapies once they have been developed. In this sense, there is a great need for the identification of biomarkers that provide quantitative measures of the process of post-traumatic epileptogenesis. However, while there is extensive reporting on functional changes following TBI (Iraji et al., 2015; Johnson et al., 2012; Sours et al., 2015), studies focusing specifically on the epileptogenic process following brain trauma are still scarce. In the current work, we found that early and late epileptogenesis patients have different phenotypes of brain-wide thalamic and hippocampal connectivity, as compared with each other and with the no epileptogenesis group, suggesting that acute hippocampal and thalamo-cortical network profiles are potential biomarkers for early and late epileptogenesis following a TBI. In addition, using functional connectivity data of thalamic and hippocampal networks, we could distinguish, with almost 88% precision, which patients developed late epileptogenesis after TBI from those who did not have epileptogenesis. Our model was able to correctly predict with 87% chance the absence of epileptogenesis following a TBI, and correctly predict with 85% chance the occurrence of epileptogenesis in a 2-year period following a TBI. These data thus open the door to the use of functional imaging in the acute setting as a key source of information for the prognostication of a patient’s potential for developing epileptogenesis in the long-term following TBI.

---

**Suppl. Table 1:** Detailed T1 image acquisition parameters

| Subj | Mag Field | MRI manufacturer | MRI model | Matrix size |  | Slices | Voxel Size |  |  | TR (ms) | TE (ms) | Flip Angle |
| --- | --- | --- | --- | --- | --- | --- | --- | --- | --- | --- | --- | --- |
| 1 | 3 | SIEMENS | Skyra | 256 | 256 | 192 | 1 | 1 | 1 | 2300 | 2.26 | 8 |
| 2 | 3 | SIEMENS | Skyra | 256 | 256 | 192 | 1 | 1 | 1 | 2300 | 2.26 | 8 |
| 3 | 3 | SIEMENS | Skyra | 256 | 256 | 192 | 1 | 1 | 1 | 2300 | 2.26 | 8 |
| 4 | 3 | Philips | Ingenia | 256 | 256 | 175 | 1 | 1 | 1 | 8.20 | 3.753 | 8 |
| 5 | 3 | Philips | Ingenia | 256 | 256 | 191 | 1 | 1 | 1 | 8.25 | 3.776 | 8 |
| 6 | 3 | GE | Signa | 256 | 256 | 164 | 1 | 1 | 1 | 11.06 | 4.788 | 20 |
| 7 | 3 | GE | Signa | 256 | 256 | 188 | 1 | 1 | 1 | 11.32 | 4.86 | 20 |
| 8 | 3 | GE | Signa | 256 | 256 | 196 | 1 | 1 | 1 | 11.58 | 4.924 | 20 |
| 9 | 3 | GE | Signa | 256 | 256 | 178 | 1 | 1 | 1 | 9.13 | 3.624 | 15 |
| 10 | 3 | GE | Signa | 256 | 256 | 192 | 1 | 1 | 1 | 9.14 | 3.624 | 15 |
| 11 | 3 | GE | Signa | 256 | 256 | 152 | 1 | 1 | 1 | 9.15 | 3.624 | 15 |
| 12 | 3 | GE | Signa | 256 | 256 | 190 | 1 | 1 | 1 | 9.13 | 3.624 | 15 |
| 13 | 3 | GE | Signa | 256 | 256 | 192 | 1 | 1 | 1 | 9.15 | 3.616 | 15 |
| 14 | 3 | GE | Signa | 256 | 256 | 186 | 1 | 1 | 1 | 9.12 | 3.624 | 15 |
| 15 | 3 | GE | Signa | 256 | 256 | 200 | 1 | 1 | 1 | 9.10 | 3.616 | 15 |
| 16 | 3 | SIEMENS | TrioTim | 256 | 256 | 160 | 1 | 1 | 1 | 1900 | 3.43 | 9 |
| 17 | 3 | SIEMENS | TrioTim | 256 | 256 | 160 | 1 | 1 | 1 | 1900 | 3.43 | 9 |
| 18 | 3 | SIEMENS | TrioTim | 218 | 208 | 160 | 1 | 1 | 1 | 1900 | 3.43 | 9 |
| 19 | 3 | SIEMENS | TrioTim | 254 | 207 | 160 | 1 | 1 | 1 | 1900 | 3.43 | 9 |
| 20 | 3 | SIEMENS | TrioTim | 256 | 256 | 160 | 1 | 1 | 1 | 1900 | 3.43 | 9 |
| 21 | 3 | SIEMENS | TrioTim | 208 | 217 | 160 | 1 | 1 | 1 | 1900 | 3.43 | 9 |
| 22 | 3 | SIEMENS | TrioTim | 256 | 256 | 160 | 1 | 1 | 1 | 1900 | 3.43 | 9 |
| 23 | 3 | SIEMENS | TrioTim | 256 | 256 | 160 | 1 | 1 | 1 | 1900 | 3.43 | 9 |
| 24 | 3 | SIEMENS | TrioTim | 256 | 256 | 160 | 1 | 1 | 1 | 1900 | 3.43 | 9 |
| 25 | 3 | SIEMENS | TrioTim | 256 | 256 | 160 | 1 | 1 | 1 | 1900 | 3.43 | 9 |
| 26 | 3 | SIEMENS | TrioTim | 256 | 256 | 160 | 1 | 1 | 1 | 1900 | 3.43 | 9 |
| 27 | 3 | SIEMENS | TrioTim | 256 | 256 | 160 | 1 | 1 | 1 | 1900 | 3.43 | 9 |
| 28 | 1.5 | SIEMENS | Aera | 256 | 256 | 160 | 1 | 1 | 1 | 2000 | 3.13 | 15 |
| 29 | 1.5 | SIEMENS | Aera | 256 | 256 | 192 | 1 | 1 | 1 | 2000 | 3.13 | 15 |
| 30 | 1.5 | SIEMENS | Aera | 256 | 256 | 192 | 1 | 1 | 1 | 2000 | 3.13 | 15 |
| 31 | 1.5 | SIEMENS | Aera | 256 | 256 | 176 | 1 | 1 | 1 | 2000 | 3.13 | 15 |
| 32 | 3 | SIEMENS | Verio | 256 | 256 | 176 | 1 | 1 | 1 | 1900 | 2.93 | 9 |
| 33 | 3 | SIEMENS | Verio | 256 | 256 | 176 | 1 | 1 | 1 | 1900 | 2.93 | 9 |
| 34 | 3 | SIEMENS | Skyra | 256 | 256 | 192 | 1 | 1 | 1 | 2300 | 2.26 | 8 |
| 35 | 3 | SIEMENS | Skyra | 256 | 256 | 192 | 1 | 1 | 1 | 2300 | 2.26 | 8 |
| 36 | 3 | SIEMENS | Skyra | 256 | 256 | 192 | 1 | 1 | 1 | 2300 | 2.26 | 8 |
| 37 | 3 | SIEMENS | Skyra | 256 | 256 | 192 | 1 | 1 | 1 | 2300 | 2.26 | 8 |

**Suppl. Table 2:**
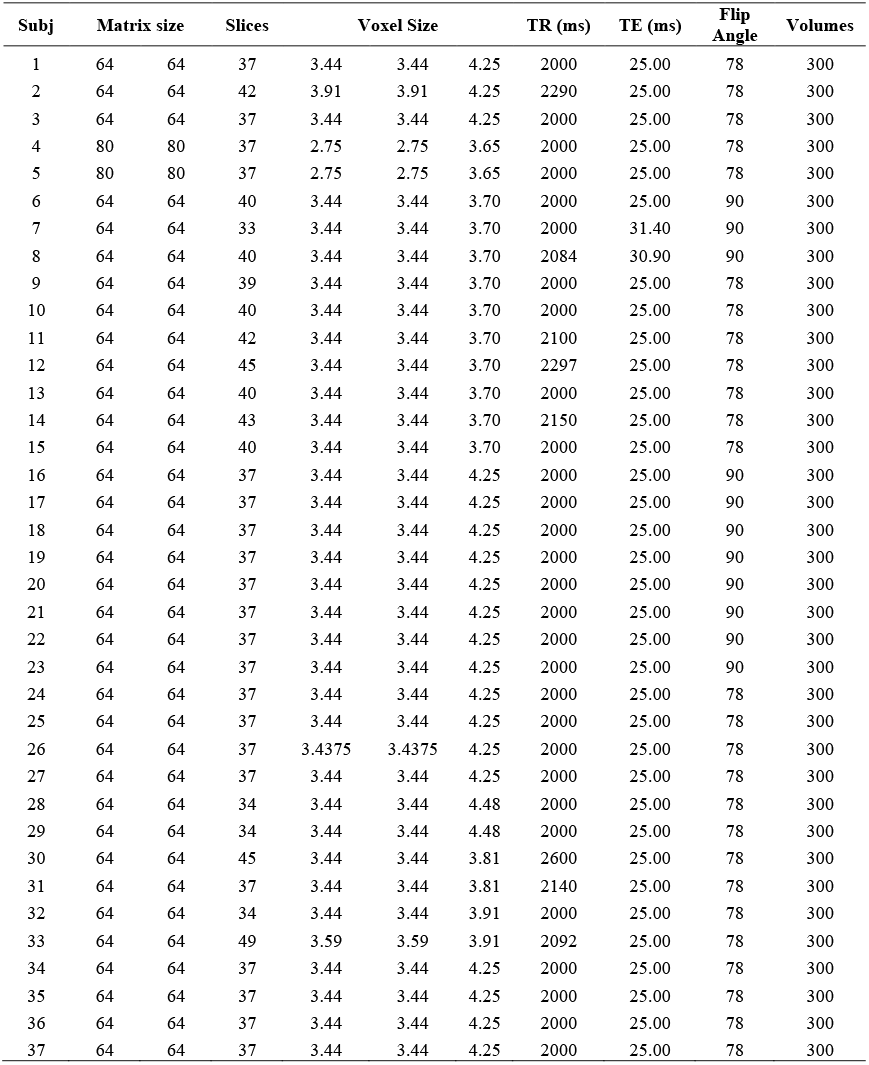
Detailed T2*-weighted echo planar images acquisition parameters

## FIGURES LEGEND

**Suppl. Fig 1:** Regions significantly correlated with the site location (*p* < .05, FWE-corrected at the cluster level), highlighting the need to harmonize the data to remove site effects on functional connectivity data.

**Suppl. Fig 2:** Distribution of thalamic functional connectivity values across sites before the harmonization (on the left) and after the harmonization (on the right). Note that the distribution curves are more similar on the right graphs, meaning that the harmonization technique diminished site effects in the functional connectivity data.

**Suppl. Fig 3:** Distribution of hippocampal functional connectivity values across sites before the harmonization (on the left) and after the harmonization (on the right). Note that the distribution curves are more similar on the right graphs, meaning that the harmonization technique diminished site effects in the functional connectivity data.

**Suppl. Fig 4**: Boxplots displaying medians, minimum, and maximum functional connectivity values for the thalamic network across sites, before (on the left) and after (on the right) harmonization of the data. Note that the functional connectivity values are more similar on the right graphs, meaning that the harmonization technique diminished site effects in the functional connectivity data.

**Suppl. Fig 5**: Boxplots displaying medians, minimum, and maximum functional connectivity values for the hippocampal network across sites, before (on the left) and after (on the right) harmonization of the data. Note that the functional connectivity values are more similar on the right graphs, meaning that the harmonization technique diminished site effects in the functional connectivity data.

**Suppl. Fig 6**: MA-plots (or Bland-Altman plots) showing the difference in thalamic functional connectivity values between two given sites for each voxel. The x-axis represents the average functional connectivity value between the two sites, and the y-axis shows the difference between them. On the left, graphs before harmonization are wider, representing larger differences in functional connectivity values between the two sites. On the right, graphs after harmonization are narrower, meaning that the harmonization diminished the difference in functional connectivity values between the two sites, removing unwanted site effects.

**Suppl. Fig 7**: MA-plots (or Bland-Altman plots) showing the difference in hippocampal functional connectivity values between two given sites for each voxel. The x-axis represents the average functional connectivity value between the two sites, and the y-axis shows the difference between them. On the left, graphs before harmonization are wider, representing larger differences in functional connectivity values between the two sites. On the right, graphs after harmonization are narrower, meaning that the harmonization diminished the difference in functional connectivity values between the two sites, removing unwanted site effects.

