## Supplementary figures and images for "Early alterations of thalamo- and hippocampo-cortical functional connectivity are biomarkers of epileptogenesis after traumatic brain injury"

### Supplemental figure 1

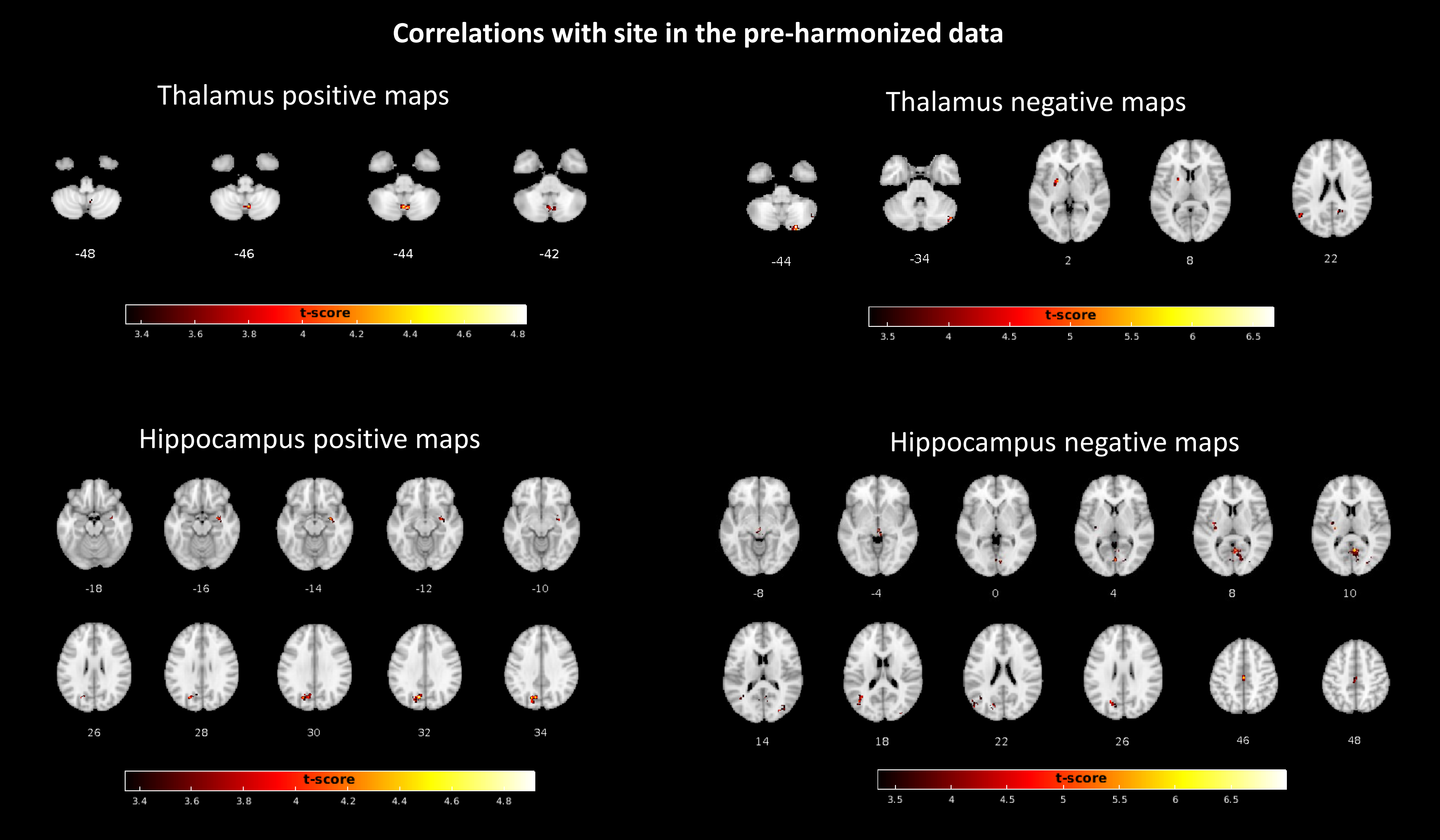

### Supplemental figure 2

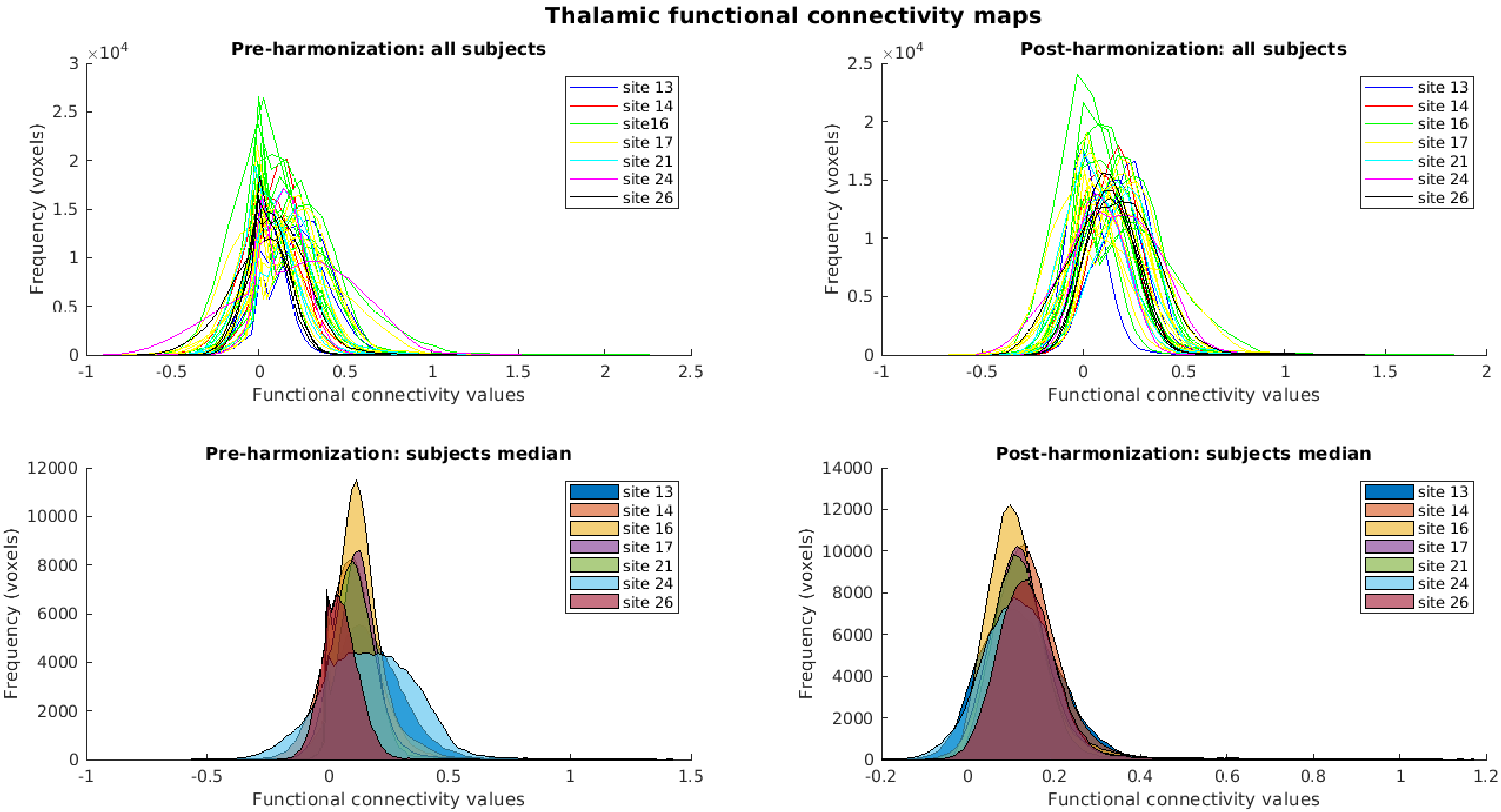

### Supplemental figure 3

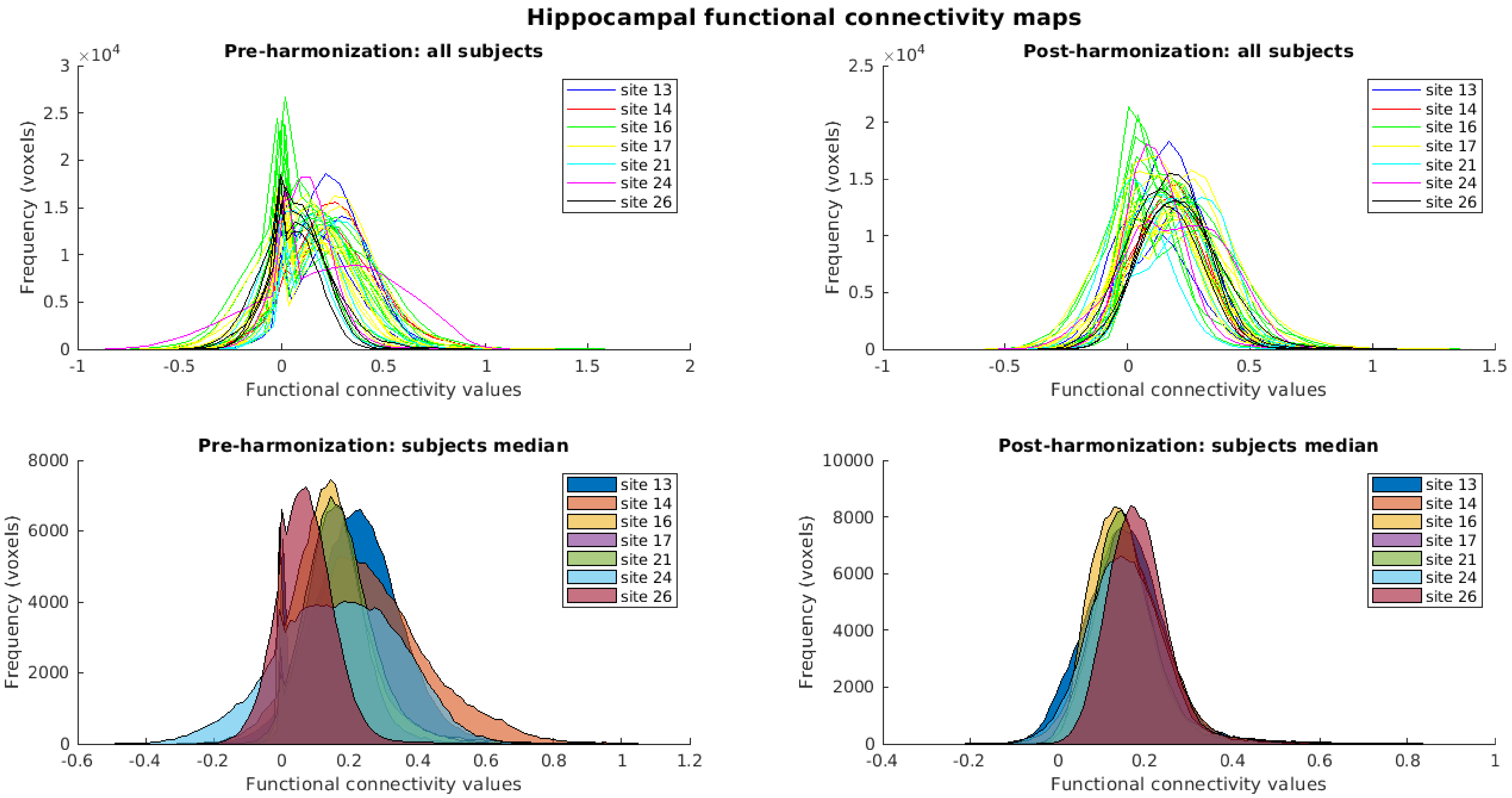

### Supplemental figure 4

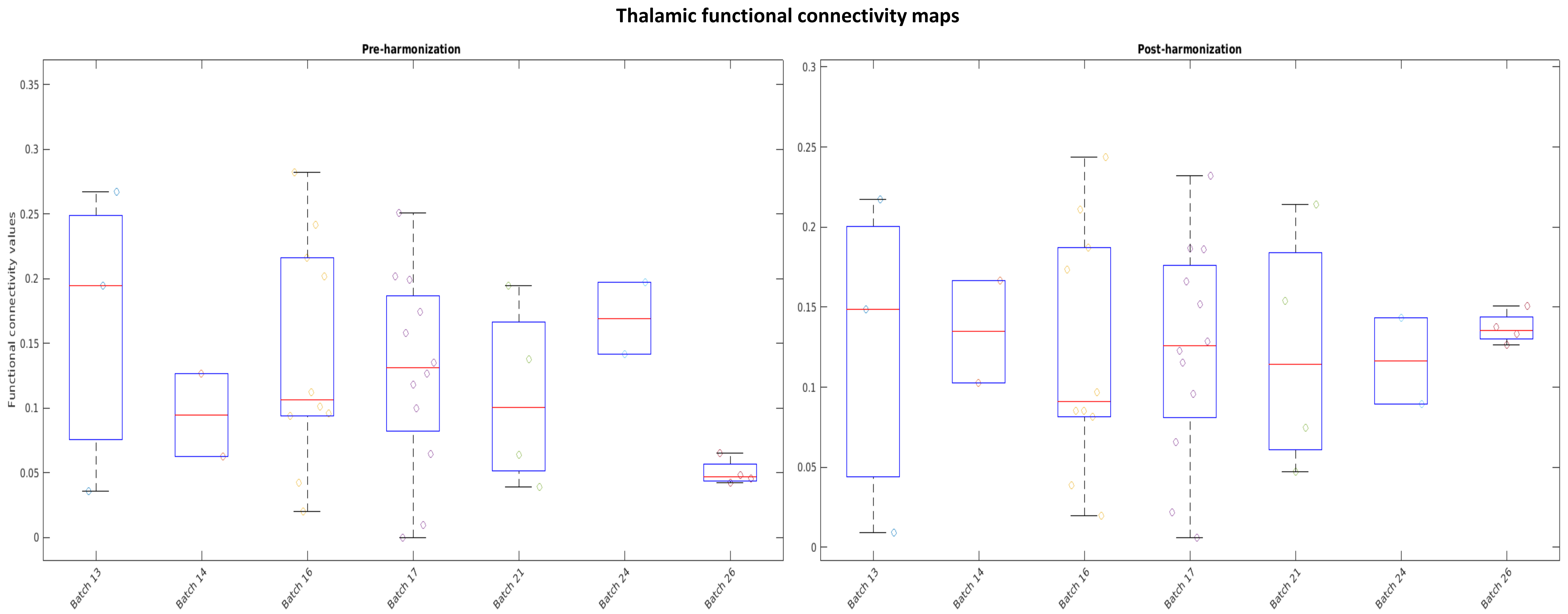

### Supplemental figure 5

#### Hippocampal functional connectivity maps

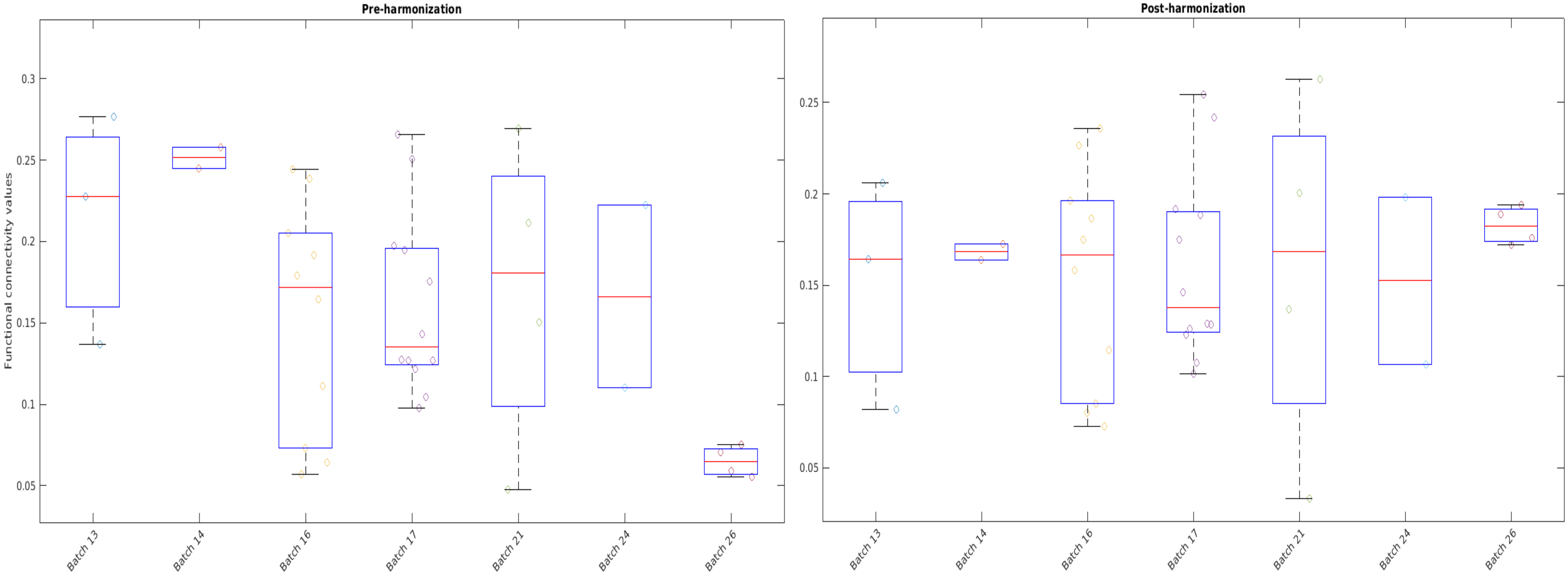

### Supplemental figure 6

#### Thalamic functional connectivity maps

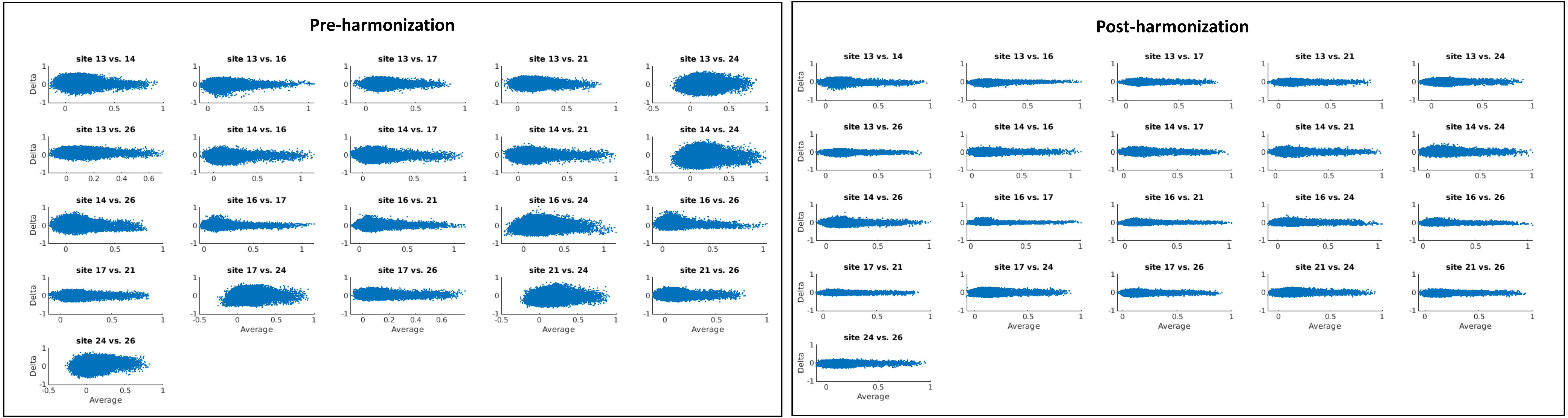

### Supplemental figure 7

#### Hippocampal functional connectivity maps

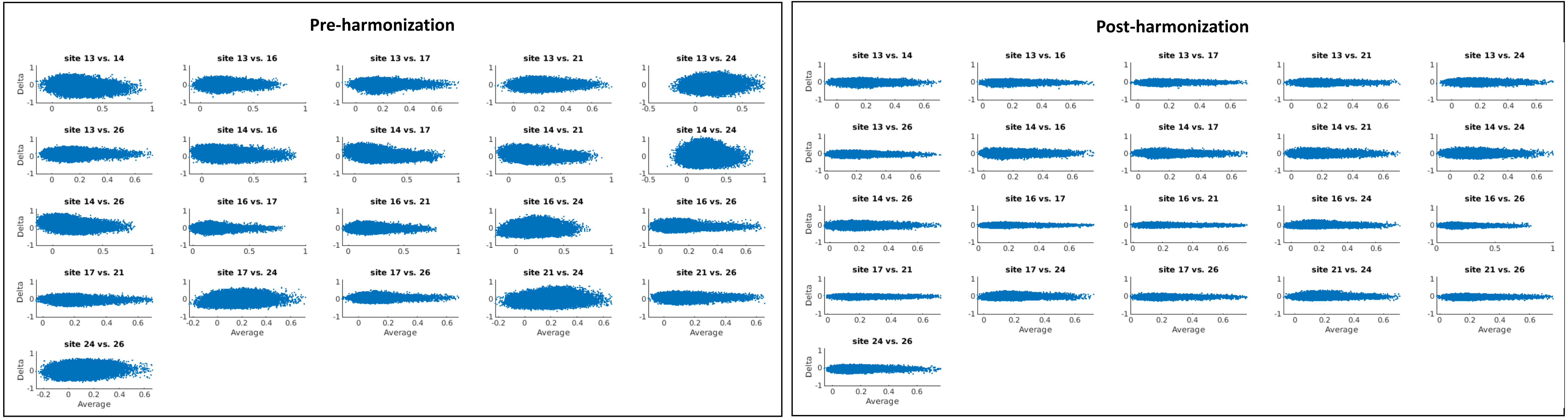
